# Mutant Plasticity Related Gene 1 (*PRG1*) acts as a potential modifier in *SCN1A* related epilepsy

**DOI:** 10.1101/282871

**Authors:** Ellen Knierim, Johannes Vogt, Michael Kintscher, Alexey Ponomarenko, Jan Baumgart, Prateep Beed, Tatiana Korotkova, Thorsten Trimbuch, Axel Panzer, Ulrich Stephani, Andrew Escayg, Holger Lerche, Robert Nitsch, Dietmar Schmitz, Markus Schuelke

## Abstract

Plasticity related gene 1 encodes a cerebral neuron-specific synaptic transmembrane protein that modulates hippocampal excitatory transmission on glutamatergic neurons. In mice, homozygous Prg1-deficiency results in juvenile epilepsy. Screening a cohort of 18 patients with infantile spasms (West syndrome), we identified one patient with a heterozygous mutation in the highly conserved third extracellular phosphatase domain (p.T299S). The functional relevance of this mutation was verified by *in-utero* electroporation of a mutant *Prg1* construct into neurons of *Prg1*-knockout embryos, and the subsequent inability of hippocampal neurons to rescue the knockout phenotype on the single cell level. Whole exome sequencing revealed the index patient to additionally harbor a novel heterozygous *SCN1A* variant (p.N541S) that was inherited from her healthy mother. Only the affected child carried both heterozygous *PRG1* and *SCN1A* mutations. The aggravating effect of *Prg1*-haploinsufficiency on the epileptic phenotype was verified using the kainate-model of epilepsy. Double heterozygous *Prg1*^-/+^|*Scn1a^wt/p.R1648H^*mice exhibited higher seizure susceptibility than either wildtype, *Prg1*^-/+^, or *Scn1a^wt/p.R1648H^* littermates. Our study provides evidence that *PRG1*-mutations have a potential modifying influence on *SCN1A*-related epilepsy in humans.

## INTRODUCTION

Epilepsy is one of the most common neurological disorders in humans, which across North America and Europe affects approximately five people in every 1000 (Banerjee *et al*, 2009). More than 350 epilepsy-associated genes have been described in the literature. Most of them play an important role in neuronal excitability, cortical development, or synaptic transmission (Noebels, 2017). The first discovered disease genes to be linked to epilepsy in humans and mice were all subunits of voltage- and ligand-gated ion channels. Mutations in these genes currently constitute approximately one third of nearly 150 monogenic seizure disorders (Noebels, 2017), affecting either voltage-gated [*SCN1A* (Escayg *et al*, 2000), *KCNQ2* (Singh *et al*, 1998)] or ligand-gated ion channels [*CHRNA4* (Steinlein *et al*, 1995), *GABRG2* (Baulac *et al*, 2001; Wallace *et al*, 2001)]. The concept of “channelopathy” implies that dysfunction of neuronal ion channels might lead to altered ion currents and destabilization of the membrane potential, potentially leading to increased epileptic network activity.

Mutations in *SCN1A* mainly cause two epilepsy syndromes, **(i)** a severe form of epilepsy characterized by fever-associated and afebrile seizures, called “Dravet syndrome” (Depienne *et al*, 2010) and **(ii)** a milder dominant familial epilepsy syndrome, called “Genetic Epilepsy with Febrile Seizures Plus” (GEFS+) (Escayg & Goldin, 2010). The severity of GEFS+ spans a broad phenotypical spectrum ranging from healthy carriers to simple febrile seizures, febrile seizures plus, and sometimes severe forms of epilepsy. On rare occasions *SCN1A* mutations may cause “Myoclonic-astatic epilepsy”, “Infantile spasms” (West syndrome), or Familial Hemiplegic Migraine (FHM) (Dichgans *et al*, 2005; Oyrer *et al*, 2018).

The genetic background, e.g. the interplay of genes jointly contributing to a biologic function such as synthesizing a protein or establishing a neuronal network, may have a profound influence on the penetrance and severity of symptoms of genetic disorders such as epilepsy. Partially, these phenomena can be modeled in mouse strains with different seizure susceptibilities. As an example, the seizure phenotype of *Scn1a* dysfunction heavily depends on the genetic background, e.g. the same *Scn1a* mutation on the 129/SvJ background results in a much milder seizure phenotype than if expressed on the C57BL/6 background (Yu *et al*, 2006). Secondly, alleles that cause mild or no phenotypes in isolation may result in more severe epilepsy when combined, as demonstrated in double mutant mice carrying the *Scn2a^Q54^*transgene together with either heterozygous *Kcnq2^p.V182M^* or *Kcnq2*^del^ (*Szt1*) alleles (Kearney *et al*, 2006). Even though the importance of various genetic factors is evident in theory, they are mostly unknown.

In Dravet syndrome, a modifying effector has been suggested that may explain the variable expressivity and penetrance of epilepsy in patients with sodium channel mutations (Singh *et al*, 2009). Different modifier genes in neural hyperexcitability pathways have been demonstrated in experimental models, e.g. comprising mutations in subunits of voltage- or ligand gated ion channels (Calhoun *et al*, 2017; Frankel *et al*, 2014). Others like the Tau protein play a general role in regulating intrinsic neuronal network hyperexcitability, and deletion of its coding gene suppresses seizures and sudden unexpected death (SUDEP) in different mouse models (Holth *et al*, 2013; Gheyara *et al*, 2014).

Beyond epileptic encephalopathies that are caused by ion channel dysfunction, epilepsy is also caused by mutations in genes involved in pathways regulating synaptic transmission (Appenzeller *et al*, 2014), especially through impairment of genes that are involved in pathways of synaptic inhibitory transmission from early development through maturation of adult GABA neurotransmission (Noebels, 2015).

Here we show that one such pathway is connected with the Plasticity Related Gene 1 (*PRG1*, syn. *PLPPR4* MIM*607813). This cerebral neuron-specific membrane protein is related to lipid-phosphate phosphatases (LPP) and is highly conserved in vertebrates. PRG1 is located at the postsynaptic density of excitatory synapses of glutamatergic cortical neurons. Postsynaptic PRG1 controls lysophosphatidic acid (LPA) signaling at glutamatergic synapses *via* presynaptic LPA2 receptors (Trimbuch *et al*, 2009) thereby reducing glutamate release probability and regulating cortical excitability from early postnatal stages (Vogt *et al*, 2017). PRG1 also affects spine density and synaptic plasticity in a cell-autonomous fashion *via* activation of the protein phosphatase 2A (PP2A)/ITGB1 (Liu *et al*, 2016, 1). To test the contribution of PRG1-deficiency to the pathophysiology of epilepsy, we investigated seizure activity in genetically modified mice after kainate application, screened human patients with West syndrome for mutations in *PRG1*, and functionally validated a mutation by *ex vivo* electrophysiology recordings in an *in-utero* electroporation model.

## RESULTS

### PATIENT STUDIES

#### Case history

The female patient (Fig. 1A, III:2) is the second child of non-consanguineous Caucasian parents. She was born at term and developed normally until the age of 6 months, when she exhibited clusters of flexion spasms and developmental regression. The EEG showed hypsarrhythmia suggestive of West syndrome. Cranial MRI and metabolic testing for increased excretion of organic acids or amino acids were normal. Seizures stopped under treatment with sulthiame. The elder brother (III:1) and both parents (II:6, II:7) are healthy. Despite her initial developmental delay, once her seizures were controlled she progressed normally and was able to achieve age-appropriate milestones later in life. Following termination of AED treatment at 2 years of age no further seizures occurred.

**Figure 1:**
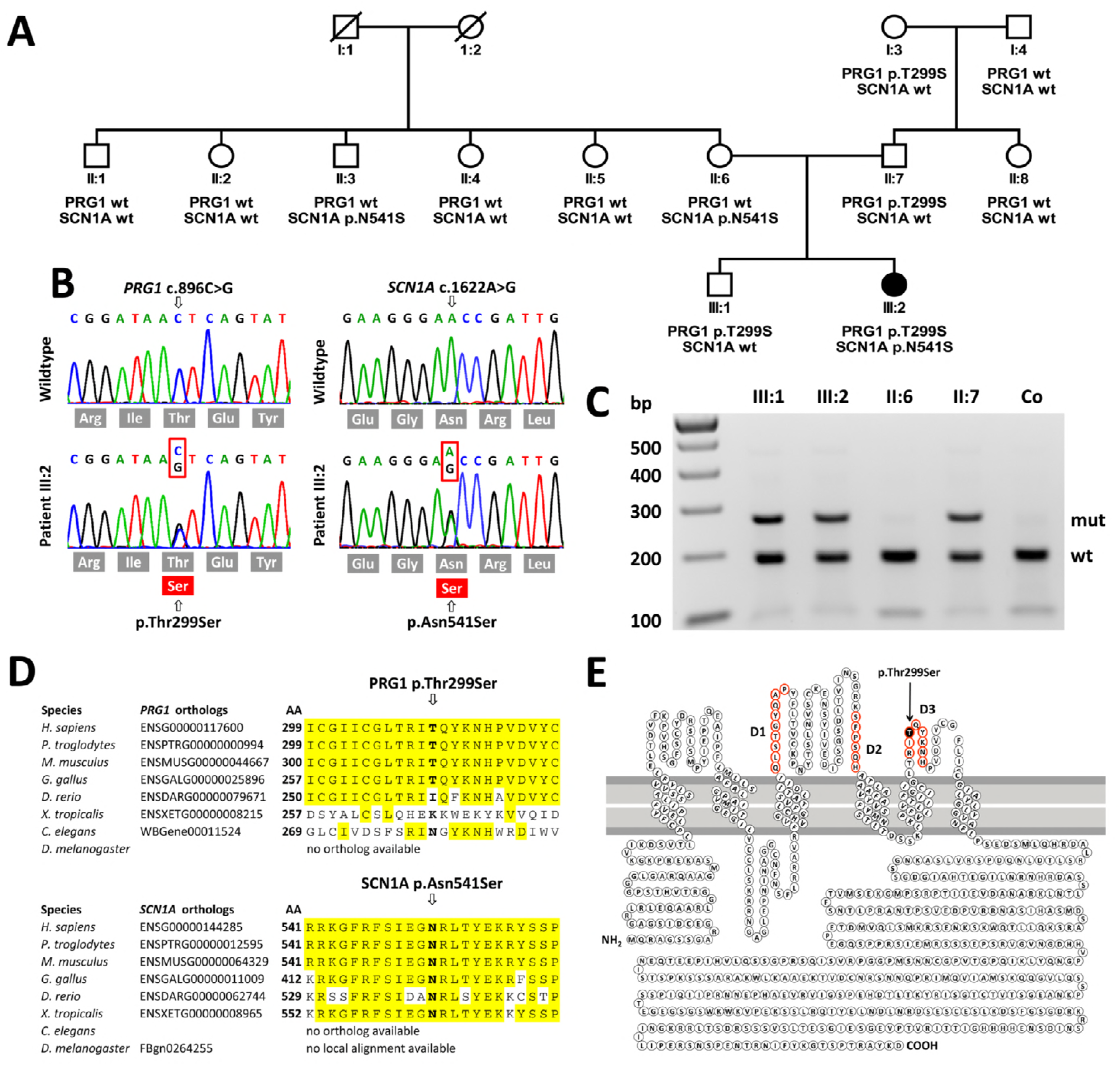
Pedigree of the family and molecular findings. **A** Pedigree of the three generation family in whom one child (III:2) was affected with West syndrome. The genotypes with respect to the *SCNA1* and *PRG1*-mutations are provided below the symbols. **B** Sequence electropherograms for *SCNA1* and *PRG1*-mutations in the index patient (lower panel) and a control (upper panel). **C** Verification of the *PRG1*-mutation by RFLP analysis where the c.896C>G mutation abolishes a *Dde*I endonuclease restriction site. **D** The multiple species amino acid sequence alignments of PRG1 and of SCN1A demonstrate the evolutionary conservation of both residues and their neighboring amino acids. The mutated amino acids are highlighted. **E** Putative structure with 6 transmembrane domains and 3 extracellular loops. The specific domains for the interaction of lipid phosphate phosphatases with lipid phosphates are highlighted in red (D1-D3). The p.T299S mutation is located in the specific D3 domain of PRG1.

#### Genetic screening revealed a combined heterozygous mutation in the *PRG1* and the *SCN1A* **gene**

We analyzed the entire coding and flanking intronic sequences of *PRG1* in a cohort of 18 unrelated patients with idiopathic infantile seizures. In patient III:2 we identified a heterozygous missense mutation [chr1:99.767.383C>G (hg19), c.896C>G, p.T299S, NM_014839] in exon 6 (Fig. 1B, C). The mutation is located in the third extracellular domain (Fig. 1E), which is evolutionary conserved in mammals and birds as well as in other LPP protein-family members (Fig. 1D). The c.896C>G variant was absent in 400 alleles of normal controls from Middle Europe as well as in the individuals of the 1000 genome and 5000 exome projects. It was found once in heterozygous state in one individual from Europe (Non-Finnish) amongst 245,604 alleles from the gnomAD database (http://gnomad.broadinstitute.org | accessed March 2018) (Lek *et al*, 2016). The heterozygous p.T299S variant, predicted to be ‘disease causing’ by MutationTaster2 (Schwarz *et al*, 2014) with a probability of P=0.983, had been inherited from her clinically unaffected father (II:7) letting us assume that the *PRG* variant might be a modifying factor of a preexisting mutation on another gene. Hence we investigated this possibility and screened for other epileptogenic mutations in patient III:2 by Whole-Exome Sequencing (WES).

WES revealed a second heterozygous missense variant [chr2:166.901.593T>C (hg19) c.1622A>G, p.N541S, NM_006920] in the *SCN1A* gene. The variant amino acid position is located in a sequence motif that is highly conserved in vertebrates (Fig. 1D) and was not listed either as a polymorphism or pathogenic variant in the *SCN1A* mutations databases (http://www.scn1a.info/; http://www.molgen.vib-ua.be/scn1amutations/). Further it was absent from the individuals of the 1000 genome and 5000 exome projects as well as from the 276,938 alleles of the gnomAD database.

The heterozygous p.N541S *SCN1A* variant had been inherited from her healthy mother (II:6) and was also present in one of the mother’s unaffected brothers (II:3). This allows the assumption that the mutation was inherited from the patient’s grandparents (I:1 or I:2) who were not available for genetic testing. Other potential disease mutations within a panel of genes presently known to cause epilepsy if mutated (n=350 on Supplementary table 01), especially those associated with West Syndrome could be excluded, either due to their frequency in healthy controls (of the 1000 Genome Project and the gnomAD Server) or due to an entirely distinct clinical phenotype (Supplementary table 02).

### ANIMAL STUDIES

#### Heterozygous *Prg1*-mutant mice show neither juvenile seizures nor spiking pattern in the EEG

In agreement with the previous reports on increased early postnatal neuronal network excitability (Vogt *et al*, 2017) and on juvenile hippocampal seizures (Trimbuch *et al*, 2009), *Prg1^-/-^*mutants (n=7) recorded on postnatal days 19-22 showed epileptiform activity in the cortical EEG and exhibited associated tonic-clonic or clonic motor seizures (4 of 7 mice, Fig. 2). One *Prg1^-/-^*mutant mouse died between P21-P22 in *status epilepticus*. Neither heterozygous *Prg1^+/-^* (n=7) nor wild type littermates (n=6) displayed pathological electrographic activity or spontaneous seizures (Fig. 2). Breeding observations of this mouse line for at least 24 months indicated that homozygous *Prg1^-/-^* animals, which survived the critical time period at around 3 weeks of age, remained henceforth clinically seizure-free until death.

**Figure 2:**
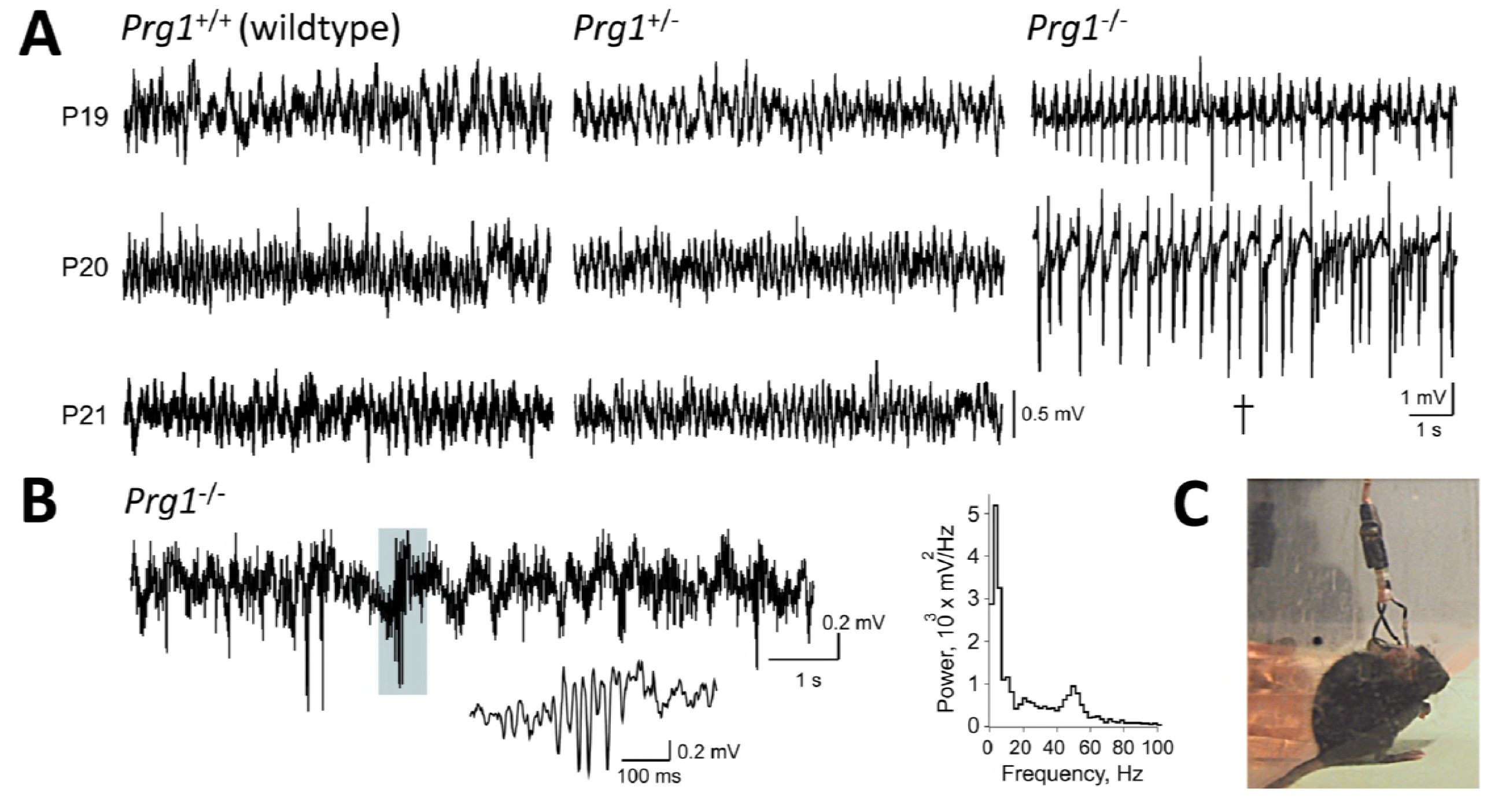
EEG recording and lack of spontaneous seizures in Prg1^+/-^ mice. **A** Examples of cortical EEG recorded in freely moving Prg1^-/-^, Prg1^+/-^, and wildtype Prg1^+/+^ littermates on postnatal days P19-21. Prg1^+/-^ and Prg1^+/+^ mice showed normal EEG, while Prg1^-/-^ mice displayed progressive aggravation of seizures up to lethal status epilepticus on day P21. **B** A hypersynchronous EEG pattern recorded on P21 in another Prg1^-/-^ mouse (left), power spectrum of this epoch (right) and a magnified slow potential with the concurrent gamma-band (∼50 Hz) oscillation (gray inset). These patterns were not observed in either Prg1^+/+^ or Prg1^+/-^ mice. **C** A snapshot from the video monitoring of motor activity performed simultaneously with EEG acquisition in a Prg1^+/-^ mouse.

#### Heterozygous *Prg1*-mutant mice show increased seizure susceptibility in adulthood

Having shown that homozygous *Prg1*^-/-^ mice had spontaneous seizures during early postnatal development, we investigated the potential of Prg1-haploinsufficiency to modify seizure susceptibility (Fig. 3). As heterozygous *Prg1*^+/-^ mice did not seize spontaneously, we used an established kainate-model in adult animals (McLin & Steward, 2006). Since the *Prg1*-knockout animals are maintained on the congenic C57BL/6J background, which is especially resistant to kainate-induced seizures, we compared the susceptibility of the mutants to their wildtype (wt) littermates. Only 7 out of 13 wt mice reached status epilepticus, which agrees with published data (McLin & Steward, 2006). After an initial kainate injection, heterozygous *Prg1*^+/-^ and homozygous *Prg1*^-/-^ mice exhibited significantly higher average seizure susceptibility scores than their wt littermates, in which epileptic activity was almost absent (Fig. 3A). In addition, *Prg1*^+/-^ mice required significantly lower amounts of kainate to progress into their first epileptic seizure than their wildtype littermates (Fig. 3B). In fact, 23 out of 24 *Prg1*^+/-^ mice and all (14 out of 14) *Prg1*^-/-^ exhibited status epilepticus, which is in strong contrast to 7 out of 13 in the wildtype group (Fig. 3C). The body weight of the mice did not significantly differ between the groups (Fig. 3D). These data suggest that Prg1-haploinsufficiency significantly increases susceptibility for entry into status epilepticus, indicating that already a partial reduction of functional Prg1 at the synapse has important functional consequences for hippocampal network stability.

**Figure 3:**
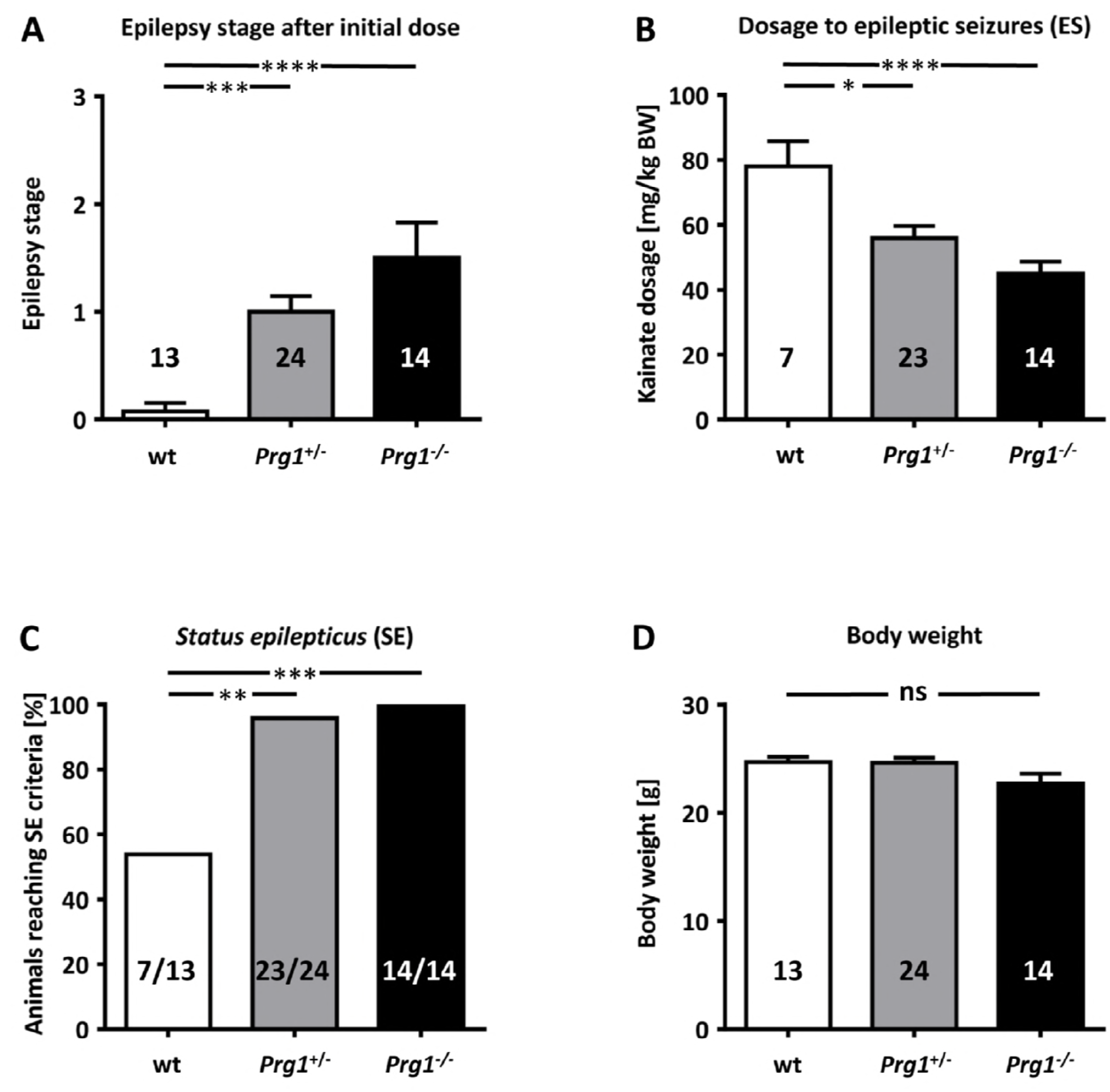
Epileptic susceptibility of adult *Prg1*^+/-^ and *Prg1*^-/-^ mice after kainate injection. *Nota bene*: as the genetic background of the mice plays an important role with regard to the seizure susceptibility upon kainate injection (McLin & Steward, 2006), the control animals in this experiment were hence always taken from wildtype littermates. **A** The epilepsy stage reached after an initial kainate dosage was significantly higher in heterozygous Prg1^+/-^ (n=24) and homozygous Prg1^-/-^ mice (n=14) than in their wildtype littermates (n=13). **B** In line with higher susceptibility to the initial dose, heterozygous Prg1^+/-^ (n=23) and homozygous Prg1^-/-^ mice (n=14) needed lower amounts of kainate to evoke epileptic seizures (stage 4). **C** Epileptic susceptibility assessed by the ability of the mice to reach a status epilepticus (SE=stage 5): only 53% of wildtype mice reached stage 5 while most (>95%) of the heterozygous and all (100%) of the homozygous *Prg1*-mutant mice reached *status epilepticus*. **D** To avoid bias by confounders, we compared the body weights between the tested groups but did not find any significant differences. Statistical analyses for panels A, B, and D were performed using the nonparametric Kruskal-Wallis with post hoc Dunn’s test, for panel C with the Pearson’s χ^2^ test. Error bars depict the SEM; significance levels: *, p<0.05; ** <.0.01; ****, p<0.001; *****, p<0.0001.

#### Double heterozygous *Prg1/Scn1a*-mutant mice show increased seizure susceptibility in adulthood

We investigated the potential of *Prg1* haploinsufficiency to modify the epileptic phenotype of heterozygous *Scn1a*^wt/p.R1648H^ mice by crossing both lines to obtain double heterozygous *Prg1*^+/-^|*Scn1a*^wt/R1648H^ mice (Fig. 4). Again, we used the established kainate-model in adult animals to induce seizures. Due to the importance of the genetic background affecting epileptic susceptibility (McLin & Steward, 2006), we analyzed wildtype littermates from the same breeding line as controls. All *Prg1*^+/-^|*Scn1a^wt/p.R1648H^* mutants (13/13) developed epileptic seizures (stage 4) after an initial kainate dose, whereas only 3 out of 21 of the *Scn1a^wt/p.R1648H^* littermates did so. Also the epilepsy stage reached after an initial kainate dosage was significantly higher in double heterozygous *Prg1*^+/-^|*Scn1a^wt/p.R1648H^* mice than in their *Scn1a^wt/p.R1648H^* or wildtype littermates suggesting higher seizure susceptibility in *Prg1*^+/-^ |*Scn1a^wt/p.R1648H^* mice. After an initial kainate injection, all but one heterozygous *Prg1*^+/-^ |*Scn1a^wt/p.R1648H^* mice directly proceeded to *status epilepticus* (SE), while *Scn1a^wt/p.R1648H^* or wildtype littermates required additional dosages to reach stage 5 criteria (SE), which is mirrored by the significant lower total amount of kainate necessary for *Prg1*^+/-^|*Scn1a^wt/p.R1648H^*mice to progress into their first seizure (Fig. 4B). The higher seizure susceptibility of *Prg1*^+/-^ |*Scn1a^wt/p.R1648H^*mice is further reflected by the fact that 100% of these mice reached SE-criteria, while SE was reached by only 76% of the *Scn1a^wt/p.R1648H^* and 71% of their wildtype littermates (Fig. 4C). No differences were observed in body weight between genotypes (Fig. 4D).

**Figure 4:**
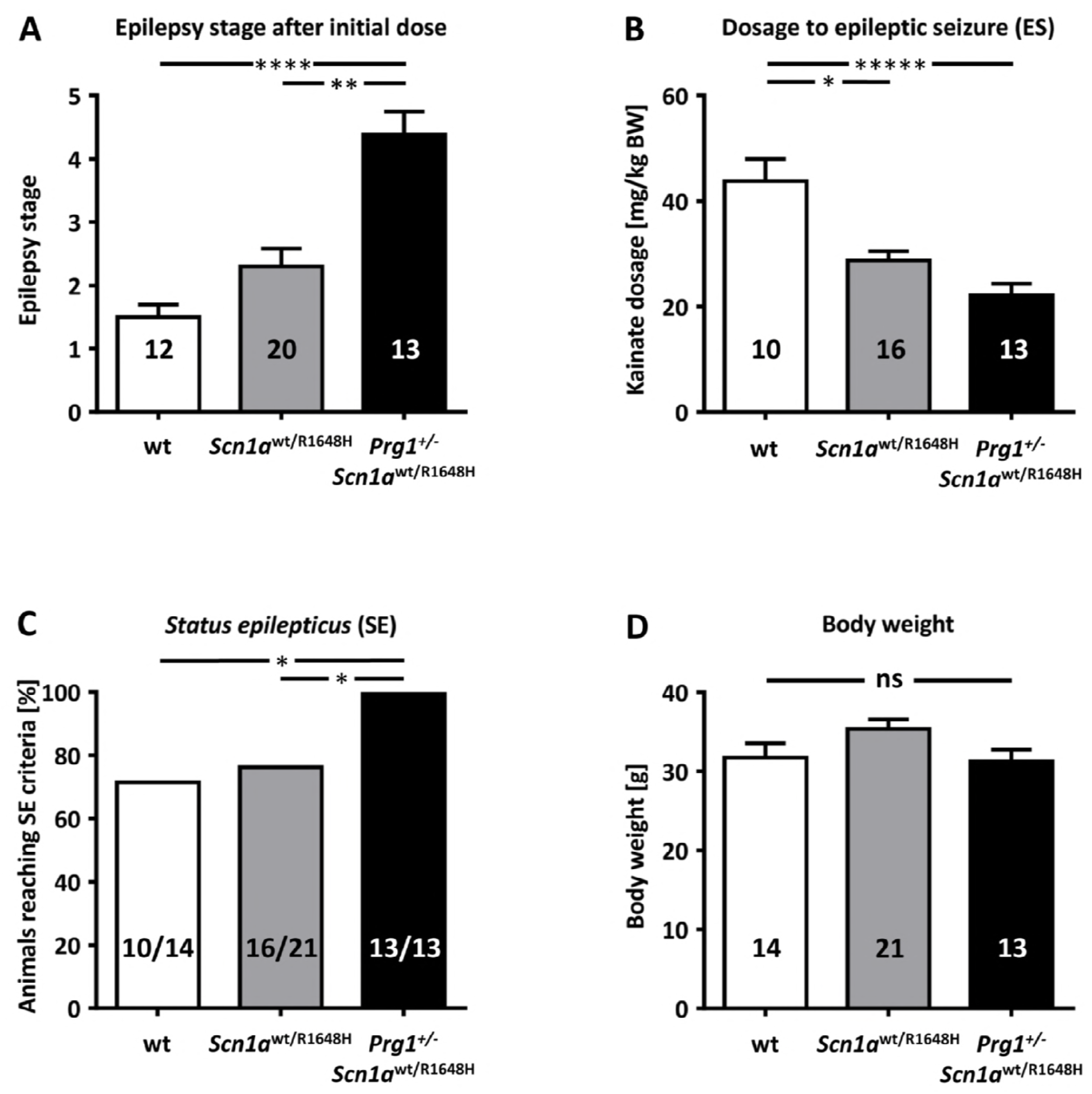
Epileptic susceptibility of adult *Scn1a^wt/p.R1648H^*, and *Prg1*^+/-^|*Scn1a^wt/p.R1648H^* double heterozygous mice after kainate injection. *Nota bene*: as the genetic background of the mice plays an important role with regard to the seizure susceptibility upon kainate injection (McLin & Steward, 2006), the control animals in this experiment were always taken from wildtype littermates. **A** The epilepsy stage reached after an initial kainate dosage was significantly higher in double heterozygous *Prg1*^+/-^|*Scn1a^wt/p.R1648H^*mice (n=13) in comparison to heterozygous *Scn1a^wt/p.R1648H^* mice (n=16) alone. **B** There was no significant difference between heterozygous *Scn1a^wt/p.R1648H^*(n=16) and heterozygous *Prg1*^+/-^|*Scn1a^wt/p.R1648H^*(n=13) mice with regard to reach epileptic seizures in response to the initial kainate dose. **C** Epileptic susceptibility assessed by the ability of the mice to reach a *status epilepticus* (SE = stage 5): only 76% of wildtype littermates reached SE while 76% of the *Scn1a^wt/p.R1648H^* and 100% of *Prg1*^+/-^|*Scn1a^wt/p.R1648H^* mice did so. **D** To avoid bias by confounders, we compared the body weights between the tested groups but did not find any significant differences. Statistical analyses for panels A, B, and D were performed using the nonparametric Kruskal-Wallis with post hoc Dunn’s test, for panel C with the Pearson’s χ^2^ test. Error bars depict the SEM; significance levels: *, p<0.05; ** <.0.01; ****, p<0.001; *****, p<0.0001.

These data suggest that *Prg1***-**haploinsufficiency significantly increases susceptibility for epileptic seizures in heterozygous *Scn1a*^wt/p.R1648H^ mice.

### ELECTROPHYSIOLOGY

#### The p.T300S mutation of Prg1 shows a loss-of-function effect in the mouse hippocampus

To test the functional relevance of the human p.T299S missense mutation of *PRG1* on the cellular level, we performed electrophysiological experiments on acute hippocampal brain slices (Fig. 5) from *Prg1*^-/-^ animals, into which we had either *in-utero* electroporated (Fig. 5A,B) a wildtype *Prg1-GFP* or a *Prg1^p.T300S^*-GFP fusion construct (*nota bene*: the p.T300S mutation in mice corresponds to the p.T299S in humans). Such functional rescue on the cellular/neuronal level has been successfully demonstrated previously (Trimbuch *et al*, 2009). This approach allowed us to investigate the electrophysiological effects of re-expressed Prg1 in a small subset of GFP^+^ single neurons independent of the surrounding *Prg1*-knockout environment.

**Figure 5:**
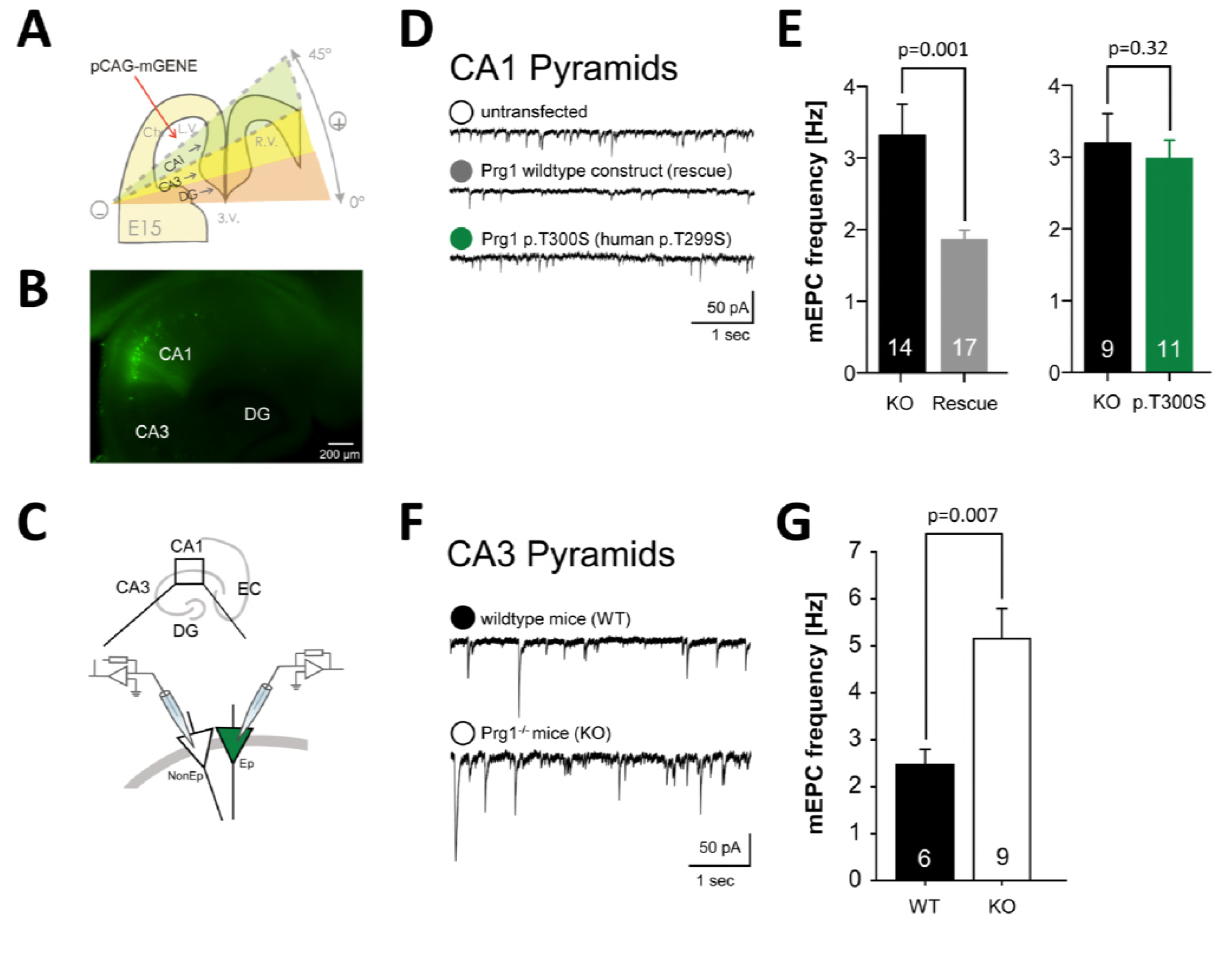
Functional testing of the Prg1 p.T300S mutation using *in-utero* electroporation and electrophysiology. **A** Scheme for *in-utero* electroporation (IUE), in which the gene of interest was introduced at E15. **B** Hippocampal slice from an IUE mouse showing the different regions. A subset of CA1 cells were successfully electroporated with the gene of interest that was GFP-tagged for visualization. **C** Recording configuration used for the miniature currents. Neighboring CA1 pyramidal cells of which one was electroporated (green, Ep) and the other non-electroporated (NonEp) were recorded. **D** Representative traces showing mEPSCs in Prg1^-/-^ (white circle), Prg1 rescued (grey circle), and Prg1^p.T300S^ (green circle) electroporated neurons. **E** Reexpression of Prg1 in Prg1^-/-^ neurons significantly decreased mEPSC when compared to neighboring Prg1^-/-^ neurons, which corresponds to a functional rescue as shown previously (Trimbuch *et al*, 2009). However, no differences in mEPSCs frequencies were observed between neighboring knockout Prg1^-/-^ neurons and Prg1^p.T300S^ electroporated CA1 pyramidal cells. mEPSC frequencies are plotted for Prg1^-/-^ neurons and *in-utero* electroporated Prg1^-/-^ neurons expressing Prg1, and Prg1^p.T300S^ respectively. The n-numbers of investigated neurons of the respective genotypes are printed on the bars. Error bars depict the SEM. **F-G** Also in hippocampal area CA3 the mEPSC frequency was significantly higher in Prg1^-/-^ mice as compared to wildtype mice. Representative traces of wildtype (black circle) and Prg1^-/-^ mice (white circle).

Whole cell patch-clamp recordings from GFP^+^/Prg1^+^ and from GFP^-^/Prg1^-/-^ CA1 pyramidal neurons in acute hippocampal slices (Fig. 5C) showed a significant decrease of the *miniature Excitatory Postsynaptic Current* (mEPSC) frequency in the GFP^+^/Prg1^+^ cells, indicating a functional rescue by electroporation of the Prg1-GFP fusion construct (GFP^-^/Prg1^-/-^, n=14; 3.32±0.43 Hz; GFP^+^/Prg1^+^, n=17; 1.86±0.13 Hz; unpaired two-tailed t-test: p=0.0014) (Fig. 5D,E).

Next we set out to test whether *in-utero* electroporation of GFP/Prg1-constructs with the mutation corresponding to human p.T299S were able to rescue the electrophysiological effects seen in the *Prg1^-/-^*neurons (Fig. 5D,E). Recordings of mEPSCs from GFP^+^/Prg1^p.T300S^ and GFP^-^/Prg1^-/-^ in CA1 pyramidal neurons did not show any significant differences in frequency (GFP^-^/Prg1^-/-^, n=9; 3.20±0.41 Hz; GFP^+^/Prg1^p.T300S,^ n=11; 2.98±0.26 Hz; p=0.32).

In the hippocampus, the neurons of the CA3 region are known to be crucially involved in epileptogenesis (Zhang *et al*, 2012). Hence we additionally performed whole cell patchclamp recordings from hippocampal CA3 pyramidal neurons (Fig. 5F**,G**) and also found an mEPSC frequency that was significantly higher in *Prg1*^-/-^ mice as compared to wildtype lit-termates (2.47±0.32 Hz versus 5.15±0.64 Hz; unpaired two-tailed t-test: p=0.007). This confirms that loss of Prg1 function increased excitability in both CA1 and CA3 areas.

These results indicate that the construct corresponding to the human p.T299S mutation was unable to rescue Prg1 deficiency on the synaptic level. The persisting increase of excitatory glutamatergic transmission functionally confirms the loss-of-function of the human mutations.

## DISCUSSION

In a previous study we show that homozygous Prg1-deficiency in mice resulted in neuronal hyperexcitability, early neuronal network synchronization, and seizures around postnatal days P18-21 (Vogt *et al*, 2017). Heterozygous *Prg1^+/-^* littermates had a normal EEG at resting state, but showed an increased susceptibility for epileptic seizures upon kainate stimulation. This indicates a sub-threshold increase of neuronal excitability caused by PRG1 haploinsufficiency.

To follow-up on these observations, we searched for *PRG1*-mutations in human patients with epilepsy. The decision, which cohorts to screen, was guided by the mouse phenotype: **(i)** Most homozygous *Prg1*^-/-^ mice convulse around postnatal days P18-21. This corresponds to a human age of 6-9 months, if referring to fundamental dynamics of brain growth, circuit organization and myelination (Levitt, 2003). The *Prg1*^-/-^ mice who survived their spontaneous *status epilepticus* lived on normally after P22 (verified by video monitoring), when seizures spontaneously ceased (Trimbuch *et al*, 2009). **(ii)** Prg1 is expressed in the mouse hippocampus during postnatal brain development (Bräuer *et al*, 2003; Unichenko *et al*, 2016), and **(iii)** the EEG of the *Prg1^-/-^* animals showed prominent irregular high-amplitude, slow-frequency discharges, multifocal spikes, and absent topical organization reminiscent of hypsarrhythmia, a hallmark of West syndrome (Dulac, 2001). We thus chose a cohort of 18 children with idiopathic West syndrome (infantile spasms).

In this cohort we found one child with a heterozygous *PRG1*-mutation resulting in the substitution of a serine for a threonine (p.T299S), which was absent in 400 Middle European control alleles, in 2,504 individuals of the 1000 genome project and present only once in heterozygous state in 122,802 individuals from the gnomAD server. Thr299 is located in the third extracellular domain in a motif that is highly evolutionary conserved in PRG1 and in other members of the LPP protein-family. Thr299 is located adjacent to Arg297, one of the critical amino acids for phospholipid interaction and de-phosphorylation of bioactive lipid-phosphates (Zhang *et al*, 2000). A similar mutation p.H253K, in the second extracellular domain that was previously introduced by *in utero* electroporation into embryonic mouse brains, disturbed the interaction between Prg1 with lysophosphatidic acid and to disrupt Prg1 function as shown by electrophysiological measurements (Trimbuch *et al*, 2009).

To establish whether the PRG1 p.T299S substitution affects protein function, we *in-utero* reexpressed the mouse homolog of the human mutation in neurons of *Prg1^-/-^* mutants and investigated glutamatergic transmission in the hippocampus. Indeed, the altered p.T299S mutant Prg1 molecules were no longer able to control lipid signaling on the synaptic level in *Prg1^-/-^* neurons. This loss of function and subsequent dramatic increase in glutamatergic transmission would likely be a contributing factor to epileptogenesis, both in mice and in humans (Bianchi *et al*, 2012).

Guided by our mouse data, where Prg1-haploinsufficiency significantly increases susceptibility for epileptic seizures, we assumed a modifying effect of the *PRG1*-variant since this variant was inherited from an unaffected parent. Our hypothesis was strengthened by the discovery of an additional p.N541S *SCN1A* variant in our patient. In our family the impact of the *SCN1A* variant alone seems to be insufficient to cause epileptic seizures as this variant had been inherited from the clinically unaffected mother and was also found in another healthy family member (Fig. 1A). This mutation, which is not present in the *SCN1A* mutation databases or in the Human Genome Mutation Database (HGMD), was predicted to be disease-causing by the MutationTaster2 software with a probability of P=0.999. It was found only once in the heterozygous state amongst 245,604 alleles from the gnomAD database, a large gene mutation databases of non-epileptic individuals. With respect to pathogenicity of a *SCN1A* variant we are aware that a number of *SCN1A* variants in the HGMD database would not be classifiable as “clearly pathogenic”, and that, as recently pointed out, a significant fraction of patients identified with *SCN1A* mutations may actually not carry any *SCN1A* variant relevant for epileptogenesis (Lal *et al*, 2016). We agree with these authors that the role of *SCN1A* missense variants in the pathogenesis of common epilepsies should not be over-stated. However, we want to point out that the pathogenicity of a certain variant does not only depend on the functional alteration of the protein in isolation, but also on the functional network in which the protein operates. In some cases this network might compensate for a minor dysfunction (as in the mother and the uncle of our patient) and in other cases not (as in our patient with West syndrome).

A number of studies provide compelling evidence for the presence of genetic modifiers. Miller *et al*. demonstrated that disease severity in *Scn1a* mutant mice strongly depends on the genetic background of the respective mouse strain and identified several modifier loci (Miller *et al*, 2014). Ohmori *et al*. reported that patients *with SCN1A* mutations plus certain *CACNA1A* variants had absence seizures more frequently than patients with *SCN1A* mutations alone and exhibited earlier seizure onset and prolonged seizure duration (Ohmori *et al*, 2013). Singh *et al*. proposed *SCN9A* as a genetic modifier of Dravet syndrome, whereby *SCN9A* may exacerbate the impact of *SCN1A* mutations on neuronal excitability (Singh *et al*, 2009). Despite these association studies, functional poof of a modifier for epilepsy in humans has not yet been provided. To study the effect of *Prg1*-mutations on a preexisting epileptic phenotype, we performed double mutant studies of *Prg1^+/-^* mice carrying an epilepsy-causing mutation in the Na_v_1.1 sodium channel (*Scn1a*). This mutation has been previously identified in a large family exhibiting either febrile or afebrile generalized tonic–clonic or absence seizures (Escayg *et al*, 2000). The *SCNA1* mouse model recapitulates the human GEFS+ phenotype, showing spontaneous generalized seizures and a reduced threshold to thermally induced seizures even in a heterozygous state (Martin *et al*, 2010). Our epilepsy studies suggest a synergistic effect with respect to seizure susceptibility in Prg1-haploinsufficient mice additionally carrying an epilepsy-causing *Scn1a* mutation, a situation which in fact resembles the genetic background of our patient harboring two mutations in a heterozygous state.

Based on the functional data, we assume that a combined haploinsuffiency of *PRG1* and *SCN1A* might be potent enough to evoke a transient severe seizure phenotype in our patient. The p.N541S variant is located in the large cytoplasmic loop between the first and second transmembrane segments of the SCN1A channel, a region known to be sometimes tolerant to substitutions, even if the substitution affects an evolutionarily conserved residue (Lal *et al*, 2016). Hence this variant might only have a modest effect on channel function, illustrated by the fact the single *SCN1A* heterozygous individuals (Fig. 1A, II:3 and II:6) are seizure free, and its epileptogenic effect only becomes manifest in combination with the heterozygous *PRG1* mutation.

In summary, our clinical and functional data indicate that PRG1-haploinsufficiency mediates an increase in excitability, sufficient to modify a pre-existing epileptic phenotype resulting in apparent aggravation and eventually seizures, but is not sufficient to cause seizures by itself. We thus assume that heterozygous *PRG1*-mutations can act as a modifier of a pre-existing epileptic phenotype. Future studies will show, whether direct modulation of PRG1 or an indirect intervention *via* the blocking of LPA2-receptors might be a valuable pharmacological tool to treat juvenile forms of epilepsy. Using pharmacological intervention into phospholipid signaling, we were able to rescue the altered cortical somatosensory filter function in an animal model with monoallelic PRG1 deficiency pointing towards a new therapeutic approach against epilepsy, e.g. *via* modulation of phospholipid signaling by pharmacological inhibition of the LPA-synthesizing molecule autotoxin (Vogt *et al*, 2016) by an orally bioavailable small molecule PF-8380 (Gierse *et al*, 2010).

### PATIENTS AND METHODS

#### Patient cohort

All patient-related studies were approved by the IRB of the Charité (EA1/215/08). All patients or caretakers provided written informed consent according to the Declaration of Helsinki. Guided by the timing of seizures and the EEG pattern of hypsarrhythmia, we selected a cohort of 18 patients suffering from idiopathic West syndrome with good outcome, who did not require antiepileptic drugs (AEDs) later in life. Brain malformations and metabolic disorders had been ruled out by appropriate imaging and metabolic studies.

#### Mutation screening in patients and control DNA samples

Genomic DNA was isolated from peripheral blood cells or from saliva by standard protocols. All coding exons of *PRG1* and 50 bp flanking intronic regions (GenBank NM_014839.4) were PCR-amplified and subjected to automatic sequencing with the BigDye® Terminator protocol (Applied Biosystems). Sequences were analyzed with the MutationSurveyor v3.10 (Soft-Genetics) and the MutationTaster software (Schwarz *et al*, 2014). PCR conditions and oligonucleotide primer sequences are available upon request. The presence of the c.896C>G *PRG1*-mutation was verified in the patient and her family by restriction fragment length polymorphism (RFLP) analysis (Fig. 1C): The oligonucleotide primer pair (FORW) 5’-TTG GCA GGC ACA GAA CAT AG-3’ and (REV) 5’-CGG CCA GAG ATT TTC TCA TT-3’ amplified a 442 bp fragment from genomic DNA, which would cleaved by *Dde*I into fragments of 190+180+72 bp in the presence of the wildtype and 262+180 bp of the mutant allele. Absence of the mutation in 200 healthy controls from the same ethnic background was verified by the same assay.

#### Whole Exome Sequencing

Exonic sequences were enriched from genomic DNA of the patient (Fig. 1A, III:2) and her parents (Fig. 1A, II:6 and II:7) using the SureSelect® V4 Human All Exon 51 Mb Kit (Agilent Technologies). Sequencing was done on a HiSeq®2500 machine (Illumina), which produced between 43-62 million 100 bp paired-end reads. The combined paired-end FASTQ files were aligned to the human GRCh37.p11 (hg19/Ensembl 72) genomic sequence using the BWA-MEM V.0.7.1 aligner (Li, 2013). The raw alignments were fine-adjusted and called for deviations from the human reference sequence (GRCh37.p11) in all exonic ±50 bp flanking regions using the Genome Analysis Toolkit (GATK v3.8) software package (DePristo *et al*, 2011; McKenna *et al*, 2010). The resulting variant (VCF) files comprised ≈60-80.000 variants and were sent to the MutationTaster2 Query Engine for assessment of the potential pathogenicity of all variants (Schwarz *et al*, 2014). Subsequent downstream analysis of potentially pathogenic variants was restricted to the 350 known epilepsy genes (Supplementary table 01). *De novo* mutations in the patient were screened for by trio-WES. Subsequently we compared the patient’s variant calling (VCF) file to those of her parents using the ‘--mendel’ option of the VCFtools v0.1.14 software package to search for variants that were present in the patient, but not in her parents (Danecek *et al*, 2011).

#### Animal studies

The animal studies were approved by the local animal welfare committee (LaGeSo T0100/03 & G0433/09, as well as G-12-096). We used male heterozygous *Prg1*^-/+^, homozygous *Prg1*^-/-^ (Trimbuch *et al*, 2009), heterozygous *Scn1a^wt/p.R1648H^* (Hedrich *et al*, 2014; Martin *et al*, 2010), and double heterozygous *Prg1*^-/+^|*Scn1a^wt/p.R1648H^*mutant mice on a C57BL/6J genetic background along with their wildtype littermates. In humans, the heterozygous p.R1648H mutation in SCN1A causes a GEFS+ phenotype (Escayg *et al*, 2000). Mice were kept under SPF-conditions with a 12 h dark/light cycle, had *ad libitum* access to food and water and were kept and euthanized in accordance with national regulations.

#### EEG recordings in freely moving animals using implanted epidural electrodes

Single tungsten wires (40 μm, California Fine Wire) were implanted into P18 pups under isoflurane anesthesia. Craniotomies were performed without damaging the underlying dura. Electrodes were placed bilaterally at 2.0 mm posterior from bregma and 3.0 mm lateral from midline with a reference electrode above the cerebellum (Trimbuch *et al*, 2009). Implanted electrodes were secured on the skull with dental acrylic. During recordings electrodes were connected to operational preamplifiers to eliminate cable movement artifacts. Electrophysiological signals were differentially amplified, band-pass filtered (1 Hz-10 kHz) and acquired continuously at 32 kHz (Neuralynx). Recordings were performed on freely moving animals at P19-P22 in 19 x 29 cm Plexiglas cages. EEG was obtained by low-pass filtering and down-sampling of the wide-band signal to 1,250 Hz. Mice were monitored from different angles by two video cameras.

#### Seizure induction with kainic acid

Adult male 3-months-old wildtype (n=13), hetero-(n=24), and homozygous (n=14) *Prg1*-mutant (Trimbuch *et al*, 2009) littermates as well as *Scn1a^wt/p.R1648H^* heterozygous (n=21) (Hedrich *et al*, 2014; Martin *et al*, 2010), *Prg1*^+/-^|*Scn1a^wt/p.R1648H^*double-heterozygous (n=13), and wildtype littermates (n=14) were analyzed for susceptibility to cerebral seizures. The susceptibility for epileptic seizures was assessed following established protocols (McLin & Steward, 2006). Briefly, animals were initially injected with 20 mg/kg kainate (at a concentration of 5 mg/ml) and assessed for 45 minutes. After this period, animals were given additional doses of kainate (14 mg/kg) at a 45 min interval (or at a 60 minutes interval after reaching level 4 seizures) and were assessed every 5 minutes. According to standard criteria from previous reports (McLin & Steward, 2006), only mice who exhibited level 5 seizures (*status epilepticus* characterized by repetitive, tonic-clonic seizures for at least 2 observation intervals longer than 10 min) during the 4 h evaluation period were included into this study.-Epileptic susceptibility (reaching stage 5) was assessed on a binary (yes/no) basis. Seizure stage was evaluated according to an established six-point scale (McLin & Steward, 2006). Epilepsy stages were evaluated by at least two independent investigators who were blinded for the genotypes. After the experiments animals tail cuts were genotyped and corresponding genotypes were assigned.

#### *In-utero* electroporation

The *in-utero* electroporation experiments in embryos from *Prg1*^+/-^ x *Prg1*^+/-^ matings were carried out in accordance with a protocol approved by the local animal welfare committee as described before (Prozorovski *et al*, 2008). The wt and mutant *Prg1-GFP* plasmids (Trimbuch *et al*, 2009) were prepared at a concentration of 4 μg/μl using the EndoFree Plasmid Kit (Qiagen). We used mice from timed matings at E15-E16 (*post coitum*). After anesthesia with 10 mg/ml ketamine and 1 mg/ml xylazine, the uterine horns were exposed. The DNA solution (1.0-1.5 μl/embryo) was injected through the uterine wall into the lateral ventricle of two of the embryos by pulled glass capillaries (WPI). Electric pulses were delivered to embryos by holding the injected brain through the uterine wall with forceps-type electrodes (CUY650P5) connected to a square-pulse generator (CUY 21 Edit, Unique Medical Imada). Five 38 V pulses of 50 ms were applied at 950 ms intervals. The uterine horns were carefully replaced into the abdominal cavity before the muscle wall and skin were sutured. Animals were checked for the *Prg1*^-/-^ phenotype after birth and the efficacy of *in-utero* electroporation was assessed by visualization of the GFP-fluorescence signal, whose coding sequence was also present on the electroporated plasmid.

#### Electrophysiology

P20-mice were anesthetized with isoflurane and decapitated. Brains were quickly removed and chilled in ice-cold, oxygenated, sucrose based artificial cerebrospinal fluid (sACSF) containing [in mM]: NaCl [87], NaHCO_3_ [26], sucrose [75], glucose [25], KCl [2·4], NaH_2_PO_4_ [1.25], MgCl_2_ [7], and CaCl_2_ [0.5] at 350±10 mOsm. Horizontal 300 μm slices were cut using a Leica VT1200 Vibratome (Leica Microsystems). Slices were then incubated for 30 min at 35°C in sACSF and afterwards stored at room temperature in normal ACSF containing [in mM]: NaCl [119], NaHCO_3_ [26], glucose [10], KCl [2.5], NaH_2_PO_4_ [1.25], MgCl_2_ [1.3] and CaCl_2_ [2.5]; pH 7.4 at 300±10 mOsm. Normal ACSF was also used for recordings. All solutions were constantly equilibrated with carbogen (95% O_2_|5% CO_2_).

Whole-cell voltage-clamp recordings were performed with an Axopatch 700B amplifier (Axon Instruments) and filtered at 2 kHz. Data were digitized (BNC-2090, National Instruments) at 5-10 kHz, recorded and analyzed with custom-made software in IGOR Pro (WaveMetrics). For whole-cell recordings, borosilicate glass electrodes (2-5 MΩ) were filled with [in mM]: K-gluconate [135], HEPES [10], Mg-ATP [2], KCl [20], EGTA [0·2], and spH was adjusted to 7.2 with KOH. Series resistance (Rs) was monitored throughout experiments; cells were rejected if Rs was >30 MΩ or varied >±30% during the recording. No Rs compensation was used. Whole-cell recordings were performed in the presence of the GABA_A_ receptor-antagonists Gabazine [1 µM] (SR 95531, Sigma-Aldrich). For the recording of miniature excitatory postsynaptic currents (mEPSCs) 2 μM Tetrodotoxin (TTX), 50 μM D-(-)-2-Amino-5-phosphonopentanoic acid (D-AP5) and 100 μM Cyclothiazide (all drugs purchased from Tocris Bioscience) were added to the recording solution.

#### Data analysis

Data was assessed for normal distribution and was analyzed accordingly. For group comparisons a nonparametric Kruskal-Wallis test was used. Non-parametric data were analyzed using the Mann-Whitney U-test. Post-hoc analysis was performed using the Dunn’s multiple comparison test. Pearson’s χ^2^ was used for dichotomous (present/absent) values. Miniature EPSCs were detected using a threshold algorithm generated in MatLab and/or Igor plug-in NeuroMatics and statistical significance was assessed with a Student’s t-test.

## ACKNOWLEDGMENTS

We thank the patient’s family for participation in this study. This work was supported by grants of the Deutsche Forschungsgemeinschaft (SFB 665 to Markus Schuelke, Robert Nitsch, Dietmar Schmitz; SFB 1080 to Robert Nitsch and Johannes Vogt, and SFB 1193 to Johannes Vogt), the European Research Council (ERC-AG “LiPsyD” to Robert Nitsch), the Stiftung Charité, and the NeuroCure Cluster of Excellence of the DFG (Exc 257) to Tatiana Korotkova, Alexey Ponomarenko, Dietmar Schmitz, Markus Schuelke.

## AUTHOR CONTRIBUTIONS

Investigated the patients, analyzed the human molecular genetics data (EK, MS); contributed to the patient cohort and patient phenotypes (AP, US); performed neurophysiologic studies (MK, AP, JB, PB, TK, TT); performed molecular genetics experiments (EK, MS, TT); performed the kainate experiments (JV, RN); contributed materials and animals (AE, HL); performed *in utero* electroporation (JV, JB); wrote the first draft of the manuscript (EK, JV, MS); jointly supervised the research (RN, DS, MS); read the final version of the manuscript and consented to its publication (all authors).

---

**Supplementary table 01:**
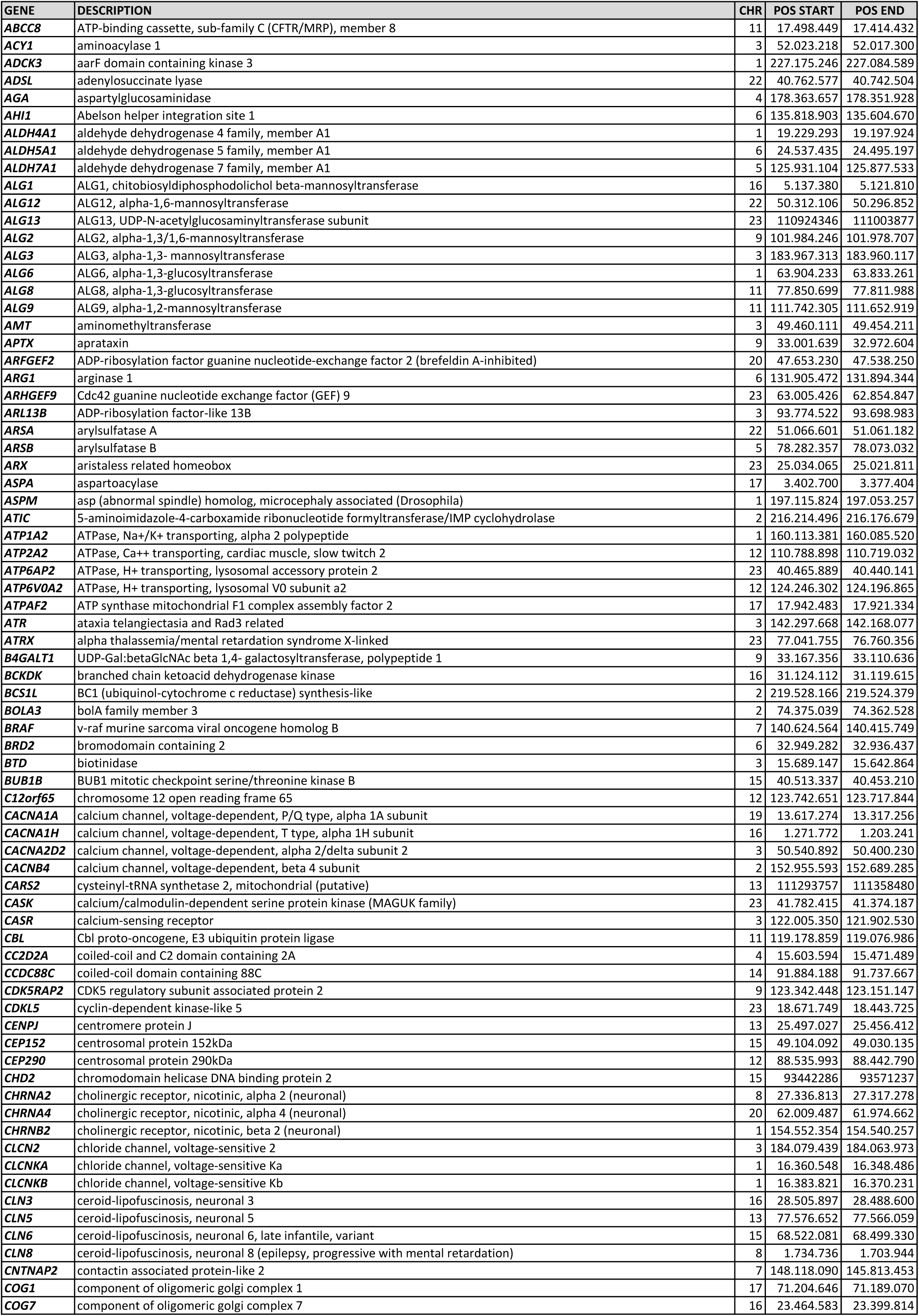
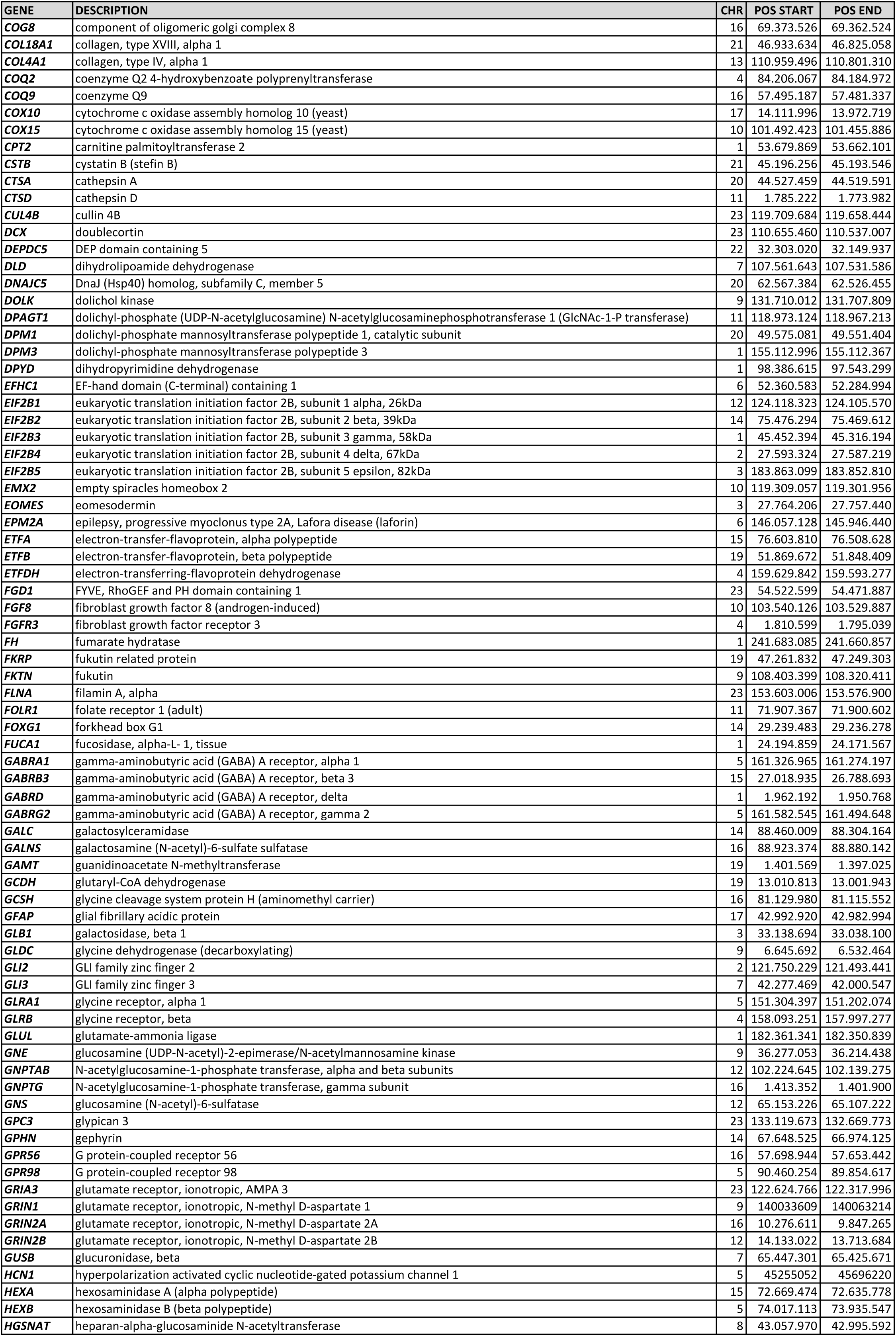
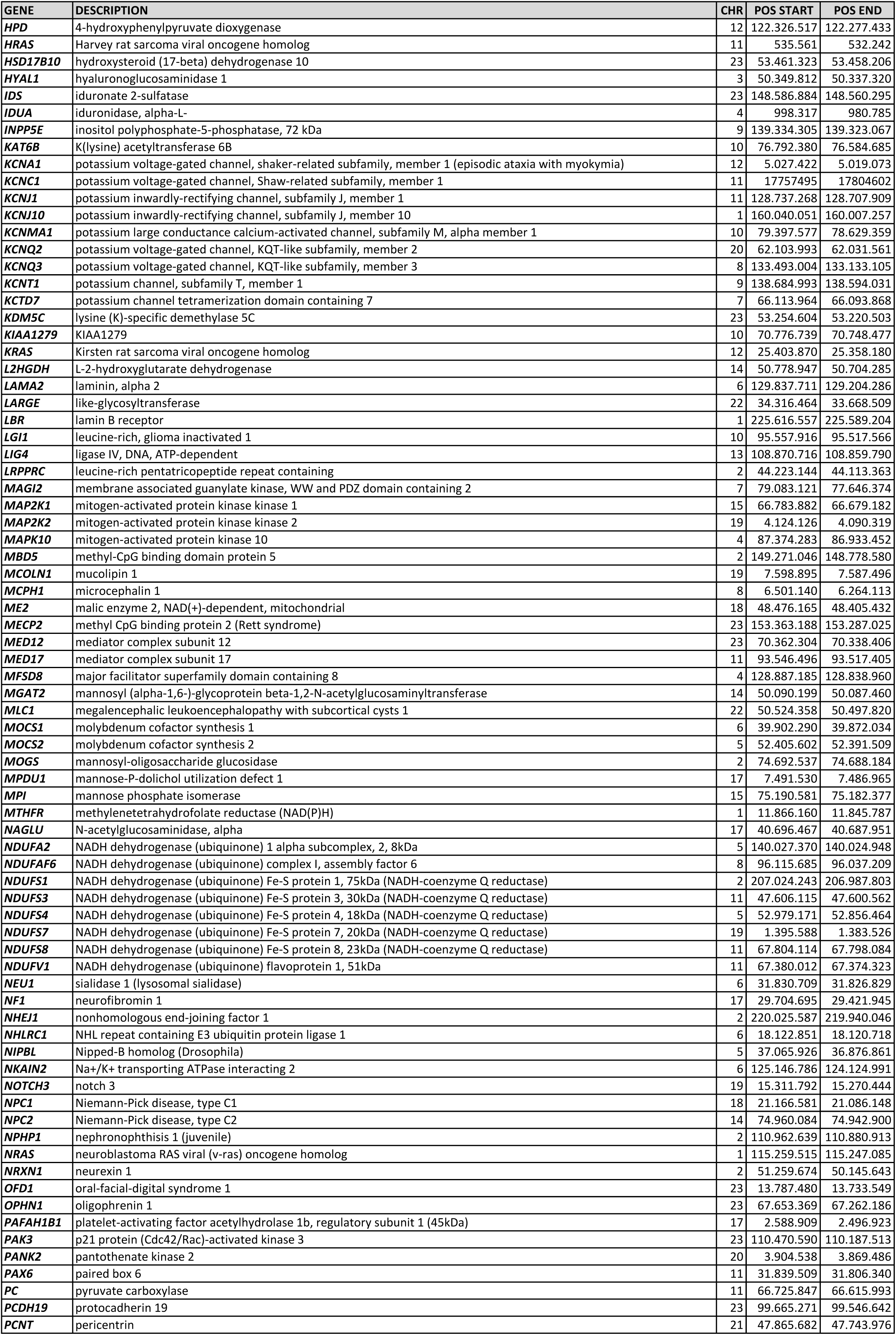
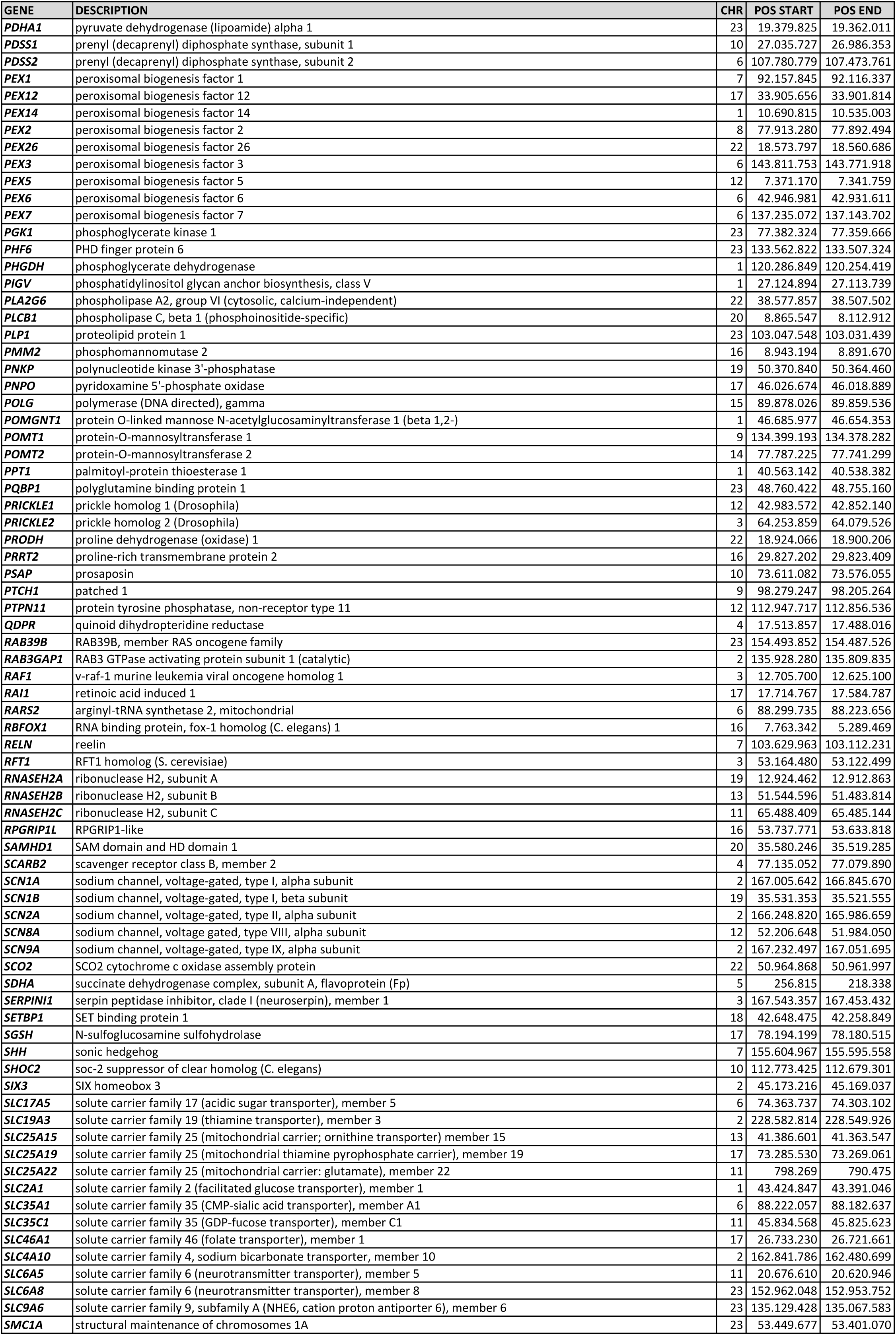
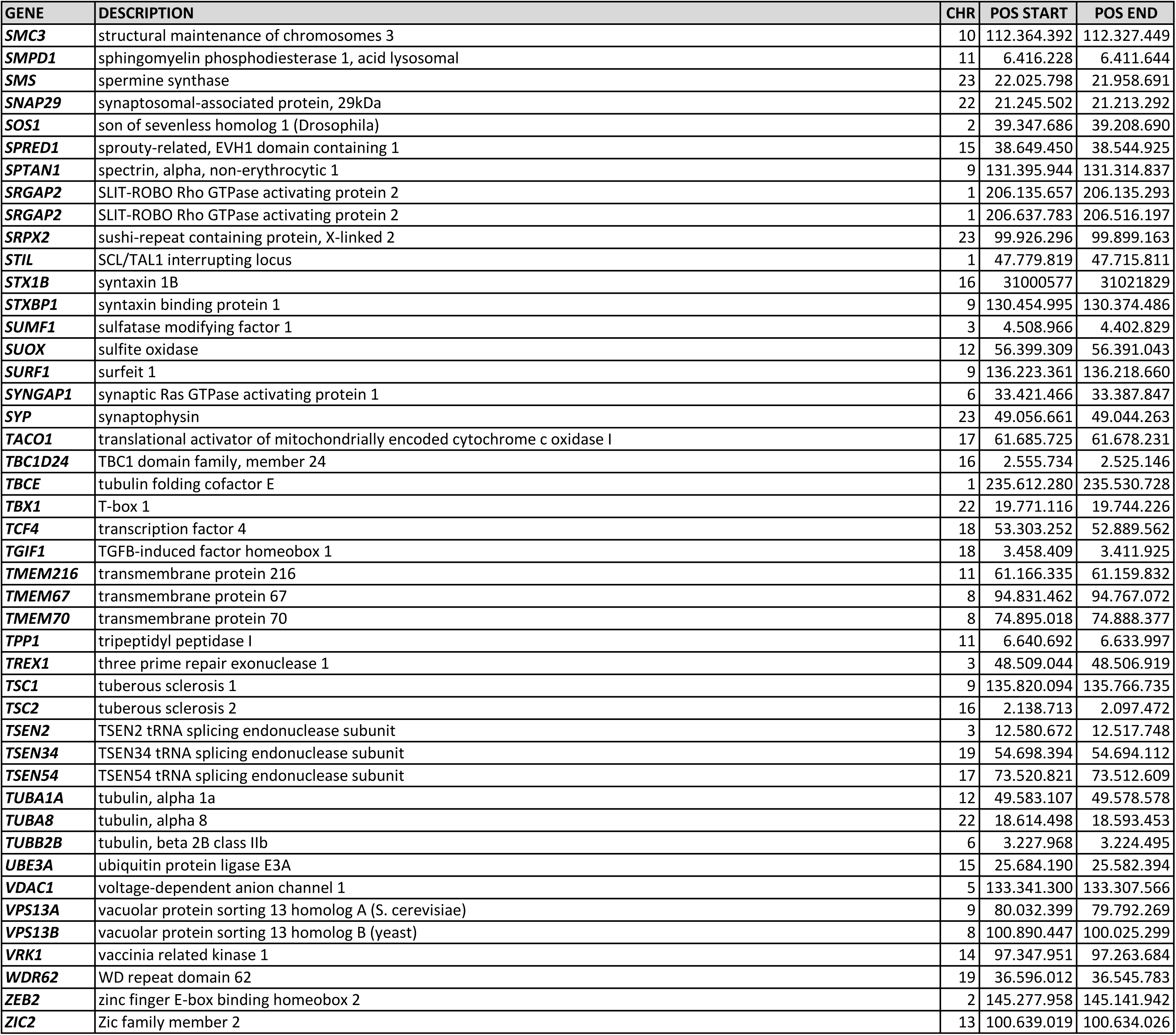
Panel of 350 genes that are known to be associated with a Mendelian trait that involves cerebral seizures (sorted in alphabetical order). The chromosomal positions refer to genome build GRCh37.p11 (hg19/Ensembl 72).

**Supplementary table 02:**
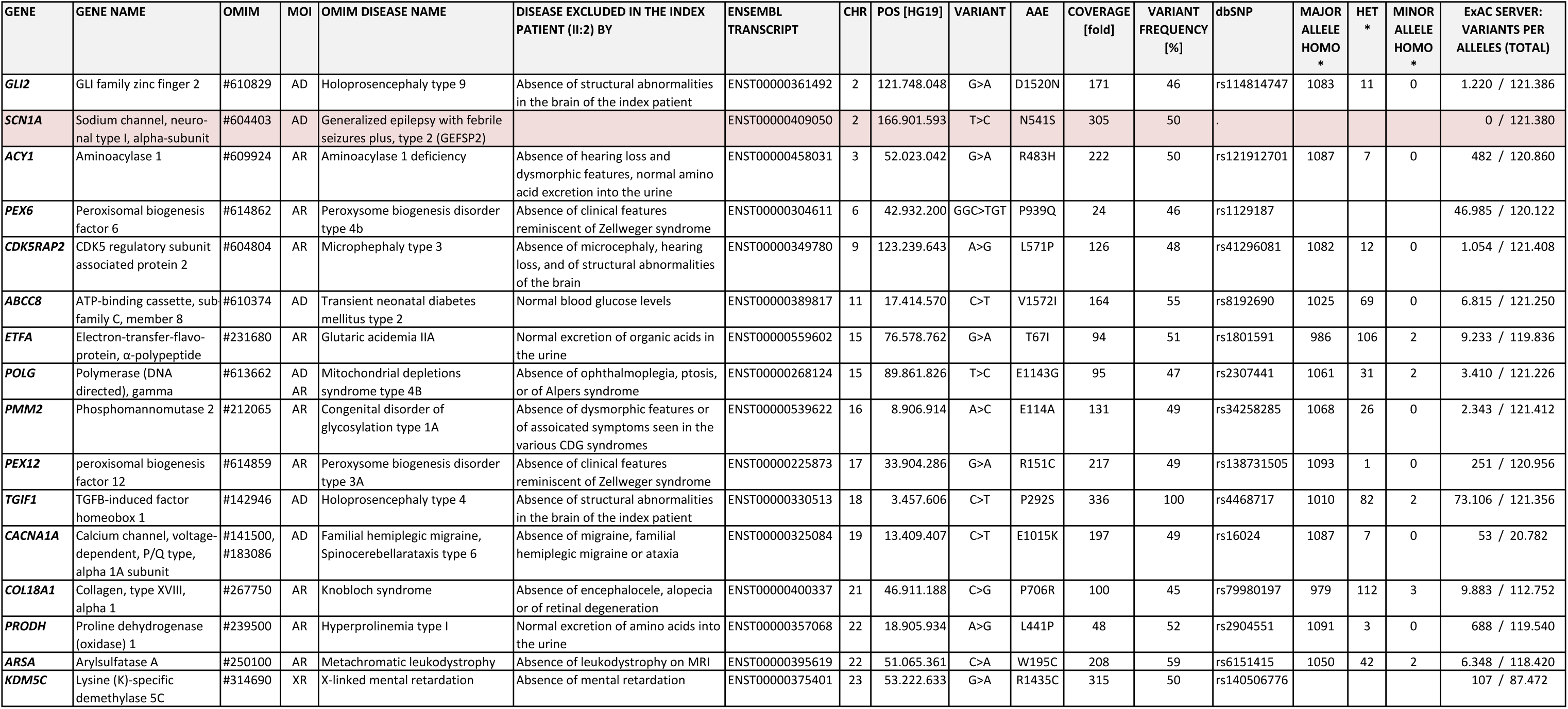
Variants causing an amino acid exchange within a panel of 343 genes associated with seizures that were predicted to be disease causing by the MutationTaster software: All variants only occurred heterozygously. Therefore diseases with autosomal recessive mode of inheritance could be ruled out. Other disorders could be excluded because the respective variant was present in the 1000 Genome project in heterozygous (HET) state or even homozygously for the minor allele or because additional symptoms characteristic for the respective disease were absent in the patient. **MOI**, mode of inheritance; **AR**, autosomal recessive; **AD**, autosomal dominant; **XR**, X-chromosomal recessive; **AAE**, amino acid exchange, * Frequencies refer to the genotypes of the 1000 Genome Project

